# The capacity to configure the hymenophore is not confined to the pore field in Oak mazegill (*Daedalea quercina*, Polyporales)

**DOI:** 10.1101/2022.08.11.503585

**Authors:** Bjarke Jensen

## Abstract

Configuring of the hymenophore of basidiocarps of Polyporales is thought to happen in the pore field immediately behind the basidiocarp margin. Basidiocarp growth is not restricted to the margin, however. Here, the importance of the pore field was assessed from observations on naturally occurring Oak mazegill (Polyporales, *Daedalea quercina*) basidiocarps and tested by experimental perturbations in natural habitats that were monitored for two years. Oak mazegill was chosen because the formed hymenophore has a unique and stable combination of poroid and lamellate features. Whether the pore field is required for basidiocarp growth was tested in ten basidiocarps in which one side was resected. New growth was observed in six basidiocarps and it occurred equally from the cut hymenophore and the intact pore field. New formation of hymenophore and pileus even occurred in seven out of ten basidiocarps that had the entire pore field resected. Whether the hymenophore is configured permanently, was tested on 54 basidiocarps on ten trunks that were turned upside-down. A new hymenophore grew through the old pileus, often far from the pore field, and its hymenophore configuration was invariably poroid despite the old hymenophore had lamellate features. In 48 experimentally banded basidiocarps, new hymenophore grew in the insertion hole of the band despite this being far from the pore field. The banded basidiocarps grew at an average rate of 5 mm per year. In conclusion, the capacity to configure the hymenophore is not confined to the pore field and it may be broadly present in the basidiocarp.

**Graphical abstract:** Graphical abstract. The configuring of the hymenophore of polypore fungi is thought to be driven by the pore field immediately behind the basidiocarp margin. The experiments on Oak mazegill basidiocarps reported here, including the complete resection of the margin and its pore field, do not support this notion. They suggest instead that the capacity to form the hymenophore is broadly present in the basidiocarp.

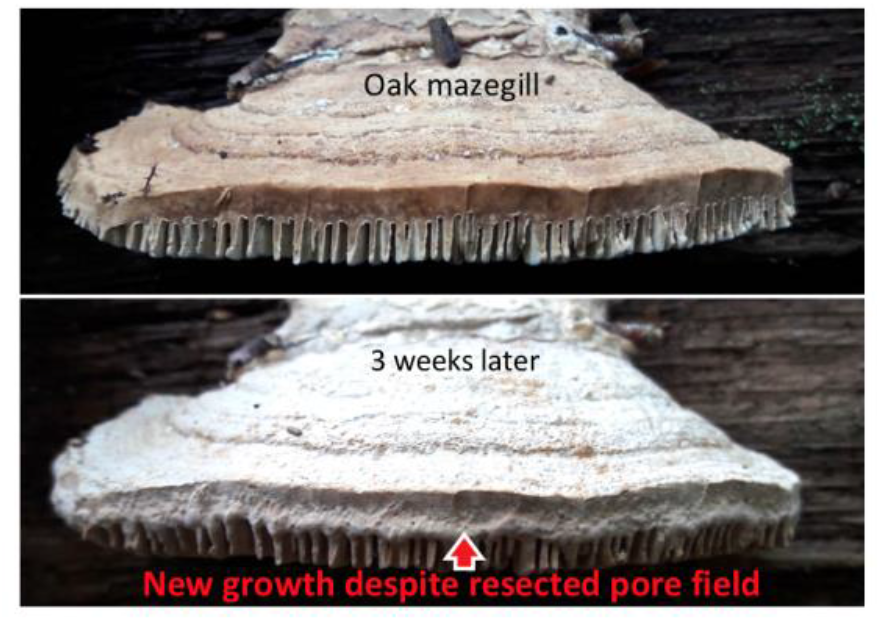

## Introduction

Carpogenesis is monophasic when all contexts of the basidiocarp are formed continuously (Clémençon 2012). Corner observed in monophasic polypores that the configuration of the hymenophore is patterned immediately behind the margin in a narrow and tube-free “pore field” (Corner 1932; Clémençon 2012). Accordingly, polypores can be considered to grow from the margin. Perennial polypores, however, will not only grow broader and wider but they will also increase in thickness, or vertical length, for example up to approximately 8 cms in the oak mazegill (*Daedalea quercina*), and the thickening is most pronounced at the base and thus furthest from the margin (Lindner et al. 2011). It is unlikely, then, that growth is limited to the margin. The margin and pore field may rather pattern the configuration of the basidiocarp. While this may be a trivial observation, it begs the question whether only the margin can configure the hymenophore. Because the hymenophore has an essential constant configuration at any part of its depth (Ames 1913 Jensen et al. 2020), it could be concluded the pore field patterns a fixed configuration.

Here, it is hypothesized that hymenophore patterning can occur outside the marginal pore field. This was tested in basidiocarps of oak mazegill by resecting the entire margin or parts thereof and by stimulating vertical growth by turning trunks with basidiocarps upside-down. Although these tests are somewhat direct tests of the importance of the pore field, one limitation to the present study is that pore field may not only be a position but it is may also be an identity to its constituent cells or context. Thus, while it is shown that hymenophore patterning can happen outside the first-established margin, it cannot be excluded that the experimentally induced patterning happens in contexts with an identity like the pore field.

## Materials and methods

### Localities

All monitored and sampled basidiocarps we found in the vicinity of the village Hald ege, Denmark, in forest in which most trees are oak (*Quercus robur).* The localities of the basidiocarps that were monitored are indicated in Figure 1 on a map from the Danish Agency for Data Supply and Efficiency (www.sdfekort.dk).

**Figure 1.**
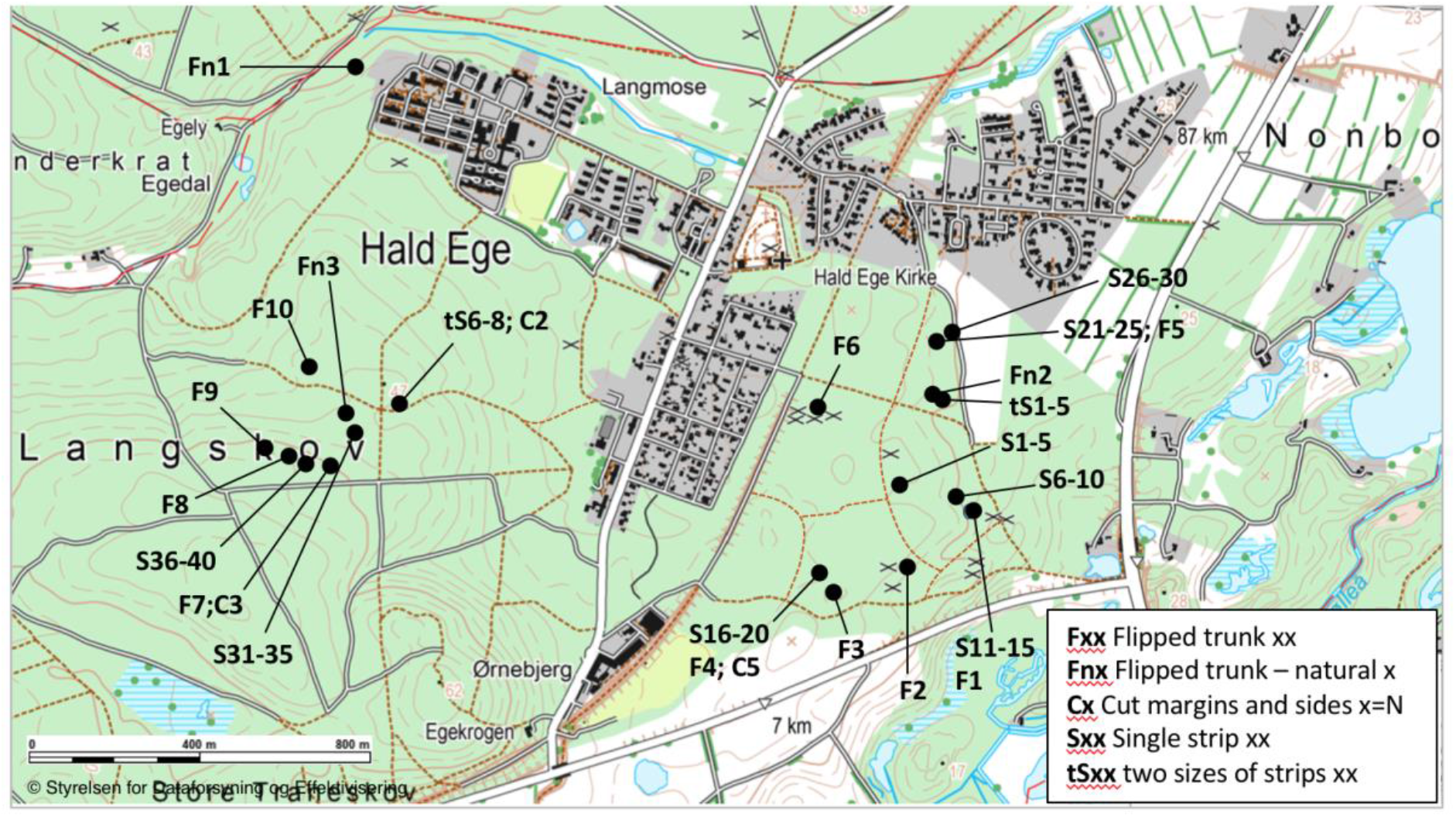
Localities at which basidiocarps were found and monitored, in the vicinity of Hald ege, Denmark.

### Growth in resected basidiocarps

To test the hypothesis that the patterning of the configuration of the Oak mazegill hymenophore is not restricted to a marginally-located pore field, we conducted two types of tests. In one test, we removed entirely the basidiocarp margin (10 basidiocarps on three trunks) and hypothesized that growth would no longer be possible. In the other test, we removed approximately 20% of the basidiocarp from one lateral side (10 basidiocarps on three trunks), and hypothesized that the basidiocarp could grow from the intact margin but not from the cut surface.

### Growth in flipped basidiocarps

We have previously observed that basidiocarps that are flipped will grow a new hymenophore through the old pileus (Jensen et al 2020 Mycologia). Here, we made observations on nine basidiocarps from three fallen trunk lying on the ground (Flipped trunk natural 1, N=5; Flipped trunk natural 2, N=3; Flipped trunk natural 3, N=1). The basidiocarps showed various signs of having grown since having been flipped by natural occurrences. On the basis of these we formulated two hypotheses that were tested by turning upside-down 10 trunks of fallen oak such that the growing basidiocarps where oriented approximately 180 degrees to their original orientation, or gravity. On these 10 trunks, a total of 121 basidiocarps were monitored for two years, numerous of which were lost or responsive. To measure the progression of the formation of a new pileus and hymenophore, we measured the area of the new hymenophore relative to the old pileus through which it was growing and the area of the new pileus relative to the old hymenophore through which it was growing. Area measurements were made using the polygon tool in ImageJ (version 1.51a, NIH, USA). The relative area of the new hymenophore was linearly correlated to the relative area of the new pileus, using a Pearson correlation test in Excel (version 16.16.27, Microsoft, USA).

### Growth around obstacles

We found 53 basidiocarps of Oak mazegill with incorporated twigs of various sizes and position in the basidiocarp (for this we did not discriminate between dead basidiocarps, which are often dark and may be overgrown with moss etc., and growing basidiocarps, which typically have a light brown pileus and an almost white hymenophore). From observations of these, we deduced how basidiocarps grow around obstacles and tested these deductions on basidiocarps in which we attached plastic strips (4.8mm wide) on five basidiocarps of various sizes on each of eight trunks (total 40 basidiocarps, 30 in Nonbo forest and 10 in Langskov). To attach the strips, a small lesion was made on the pileus surface approximately halfway between the base and the margin, either with a screwdriver or a knife, and the strip was then forced through the basidiocarp and closed such that the strip made contact with the margin. We learned that the pileus is relatively brittle in abrupt boundaries between growth zones (these are revealed as a change in color and, or, height of the pileus) and we preferentially inserted the strips through such boundaries. These basidiocarps were monitored for two years. In additional eight basidiocarps, we inserted a strip of medium size (4.8mm wide, same width as above) and a strip of large size (7.6mm wide) in each of eight basidiocarps situated on three different trunks and these were also monitored for two years.

### Estimation of growth rate

The increment in size of basidiocarps with attached plastic strips was used to deduce growth rates. Images of the basidiocarps were imported to ImageJ. Using the line tool, in each image the width of the plastic strip was measured for scale and then the outward growth was measured along a single line next to the perturbation of the margin created by the strip.

## Results

### Growth despite resected pore field

Of the 10 basidiocarps with a resected side, five exhibited pronounced growth from the cut surface already within three weeks after the resection (Figure 2A). This new growth occurred along the entire cut surface and it was not restricted to the pore field nor was it preferentially originating from there (Figure 2A). The resected parts were left next to the cut basidiocarp but they never exhibited observable growth or remodeling (Figure 2A). This suggests that the response to damage is mostly or fully dependent on nutrition from outside the basidiocarp and that the response could be considered new growth rather than reorganization. Snakes, as a contrary example, may elongate while fasting (McCue 2007). After one year, six out of 10 basidiocarps exhibited sustained growth from the cut surfaces, again showing substantial growth from the entire cut surface and no preferential growth from the pore field (Figure 2B). The configuration of the newly formed hymenophore was lamellate in some instances, showing that the poroid configuration is not a necessary configuration after perturbation (Figure 2B). The newly formed tubes were oriented perpendicular to the plane of the cut, i.e. following the direction of growth, suggesting the lamellate configuration reflects growth rather than patterning.

**Figure 2.**
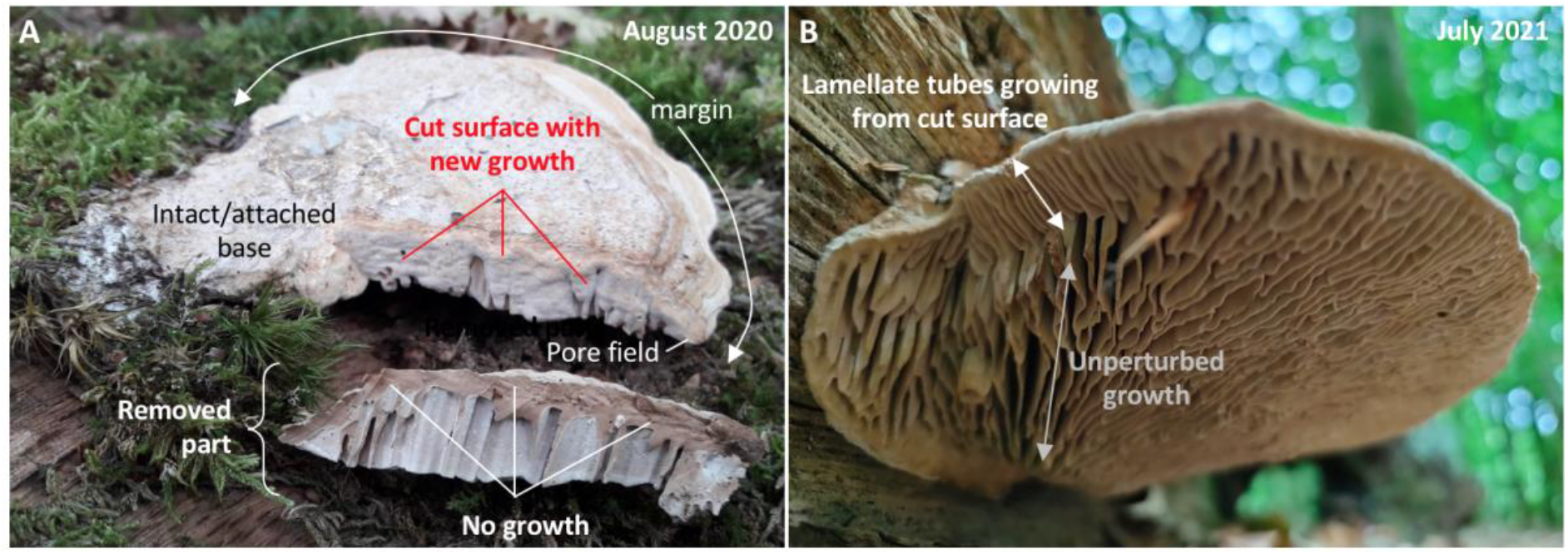
Basidiocarps with resected sides. A. Basidiocarp growing on the same trunk as ‘S16-20’. It exhibits new growth in the part that remains attached to the trunk, whereas the removed part does not show signs of new growth. The cut was made three weeks prior to the day the photos were taken (resection 26^th^ of July 2020, photography 16^th^ of August 2020). Notice that the growth response is on the entire cut surface and it is therefore not restricted to the margin where the pore field is. B. Basidiocarp growing on the same trunk as ‘tS6-8’, with a resected left side as the basidiocarp shown in A. It exhibits substantial growth from the cut surface. The new growth is not preferentially from the margin (and therefore pore field). In addition, the new hymenophore is predominantly lamellate showing that a poroid configuration is not the necessary response to perturbed growth.

Ten other basidiocarps had their entire margin, including the pore field, resected. Five of these exhibited pronounced growth from cut surfaces already within three weeks after the resection (Figure 3A-B). Two year after the resection, seven basidiocarps exhibited substantial growth of both new pileus and hymenophore from the cut surfaces (Figure 3C-E), which suggests the pore field is not required for growth. The new hymenophore that had formed distal to the cut surface was lamellate in some cases (Figure 3D). These observations suggest that growth is not dependent on the presence of a pore field. The next experiments were conceived to further test the importance of the pore field by inducing the formation of a new hymenophore while the basidiocarp margin was left intact.

**Figure 3.**
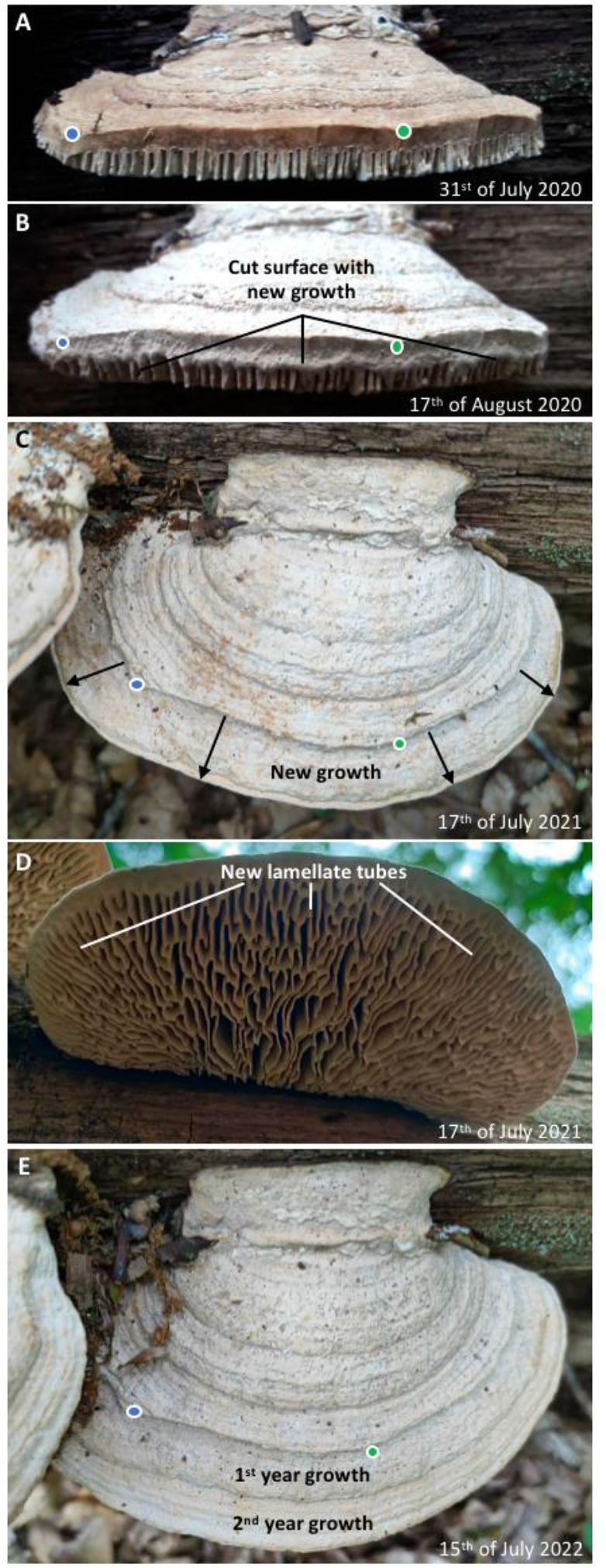
Growth despite resected margin and pore field. A. Resection of the margin of a basidiocarp from location ‘Flipped trunk 7’. Given tubes are exposed on the cut surface, the pore field must have been resected. B. An initial growth response occurred already within three weeks. C. One year after the resection, substantial outward growth had taken place (notice the same positions in A-C and E are indicated with blue and green dots). D. The hymenophore that had formed after the resection had a predominantly lamellate configuration. This shows that new growth can occur even if the pore field is resected and that a poroid configuration is not the necessary response to perturbed growth. E. After two years the basidiocarp still exhibits substantial growth.

### Hymenophore formation in flipped basidiocarps

From the following observations, it was deduced that the formation of a new hymenophore could be induced by turning basidiocarps upside-down. Nine naturally occurring basidiocarps were found with clear signs of having grown in one vertical direction and then later grown in the opposite vertical direction (Figure 4), i.e. they had been flipped by presumably natural events. These grew on three trunks from different localities (Figure 1, Flipped trunk natural 1-3). The signs of having been flipped were principally a partially covered hymenophore pattern on the present sky-facing pileus together with neighboring and darkly colored, or old, basidiocarps with sky-facing hymenophores. In the flipped basidiocarps, the new hymenophore that was vertically below the old hymenophore was characterized by a very low prevalence of lamellate tubes (Figure 4A). Lamellate tubes, however, were abundant in the part of the new hymenophore that appeared to have grown outwards since the basidiocarp was flipped (Figure 4A). Basidiocarps from the same trunk and which appeared to have started growing after the trunk was flipped showed the typical daedalean configuration with many lamellate tubes near the base (Figure 4B), as did basidiocarps from different trunks from the same location (Figure 4C).

**Figure 4.**
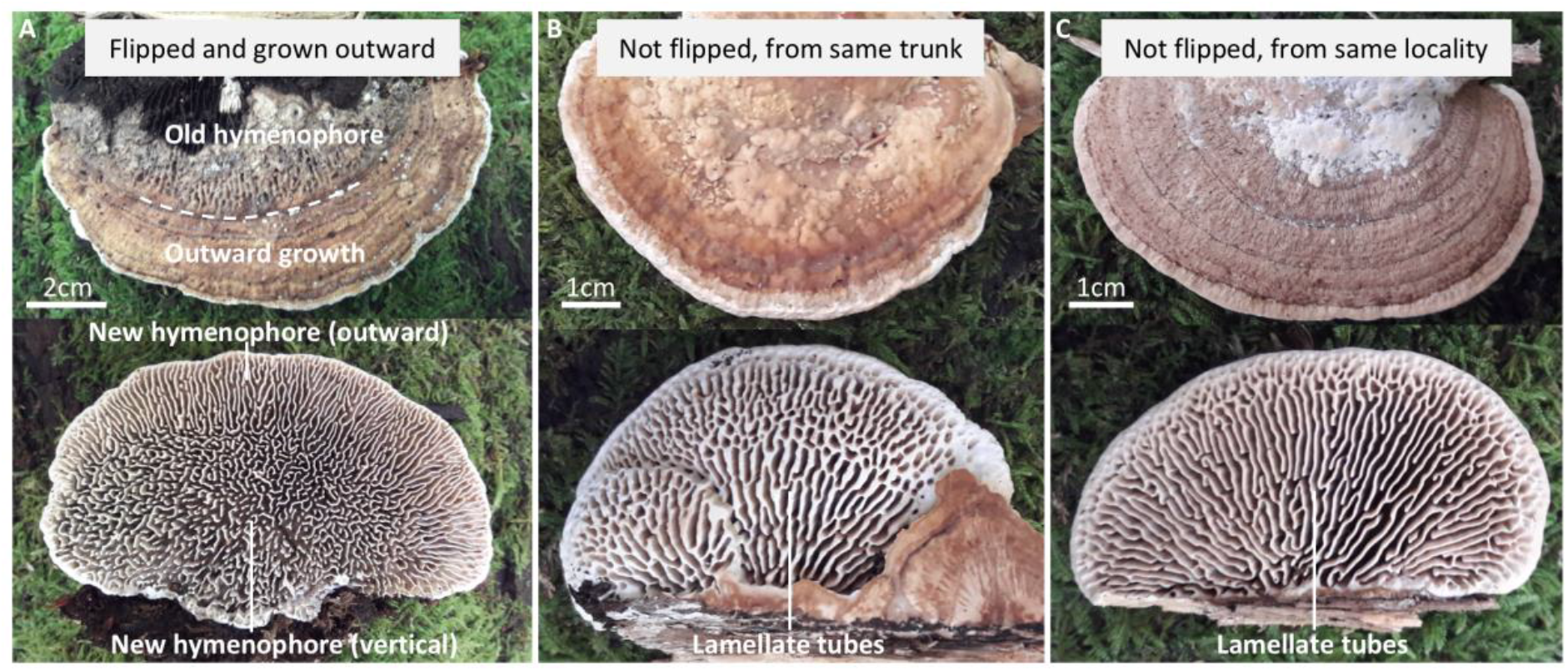
Effect of growth direction on hymenophore configuration. A. A basidiocarp flipped by natural events in which the old hymenophore is partly obscured by a new pileus and there has been substantial outward growth since being flipped. The new (vertical) hymenophore configuration is almost hydnoid and lamellate tubes with radial orientation are confined to the new part formed by outward growth. B. Basidiocarp from the same trunk as A with no obvious signs of having been flipped and in which the hymenophore configuration at the base has numerous lamellate tubes. C. Basidiocarp from the same location as A-B, but from a different trunk, in which the hymenophore configuration at the base has numerous lamellate tubes.

For the experimentally induced formation of a new hymenophore, we identified 10 trunks of oak laying on the ground and which had 121 basidiocarps growing on them. The hymenophore configuration of all basidiocarps was daedalean. The trunks were rotated approximately 180 degrees such that the basidiocarps were turned upside-down. Already three weeks after being flipped (i.e. by mid-August 2020), some basidiocarps showed clear signs of responding to the new orientation. Most noticeable was the thickening of the tubes that were now facing the sky and we found this to be occurring in 40 out of 121 basidiocarps. In a few the thickening was so advanced that pores were sealed (Figure 5A-B), while in others the thickening was in its earliest stage and was barely detectable. We also observed in three out of 121 basidiocarps the formation of new hymenophore on the old pileus that was now facing the ground (Figure 5C). In these, the configuration of the old tubes was predominately lamellate, whereas the new hymenophore was poroid (Figure 5C). Most noticeably, the new hymenophores were closer to the base than to the margin and the pore field (Figure 5C).

**Figure 5.**
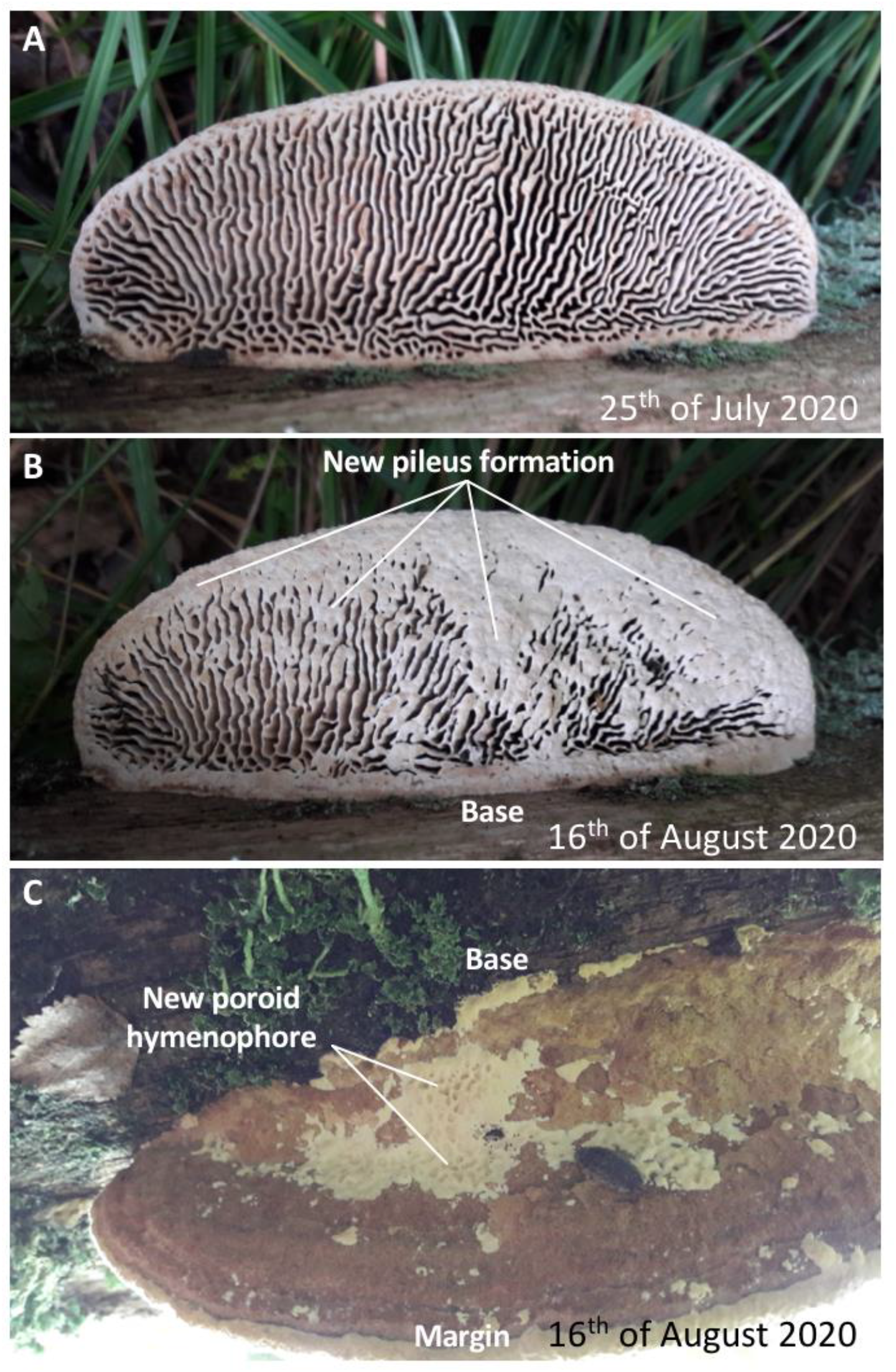
Short-term changes to experimentally flipped basidiocarps. A-B. Basidiocarp from ‘Flipped trunk 6’ where substantial thickening of the tubes has occurred in 22 days. C. New hymenophore in poroid configuration has started to develop on the pileus that faces the ground.

The same 121 flipped basidiocarps were reassessed after one year and 66 exhibited a new pileus and, or, hymenophore (Figure 6). In 44 cases a new hymenophore was forming and in ten a new hymenophore covered the entire new underside. No tubes were ever configured as lamellae in contrast to what could be expected if the hymenophore configuration is fixed. Based on the analysis of 59 basidiocarps, where either a new pileus and/or hymenophore was forming, we found the formation of a new pileus had on average proceeded further than the formation of a new hymenophore (Figure 6D) (seven out of 66 basidiocarps were excluded because of 100% coverage of the new pileus and hymenophore). When analyzed as a linear regression, there was a 0.4% increase in new hymenophore coverage for every 1% increase in new pileus coverage (P<0.001, alpha=0.40, 95% confidence interval 0.19-0.60).

**Figure 6.**
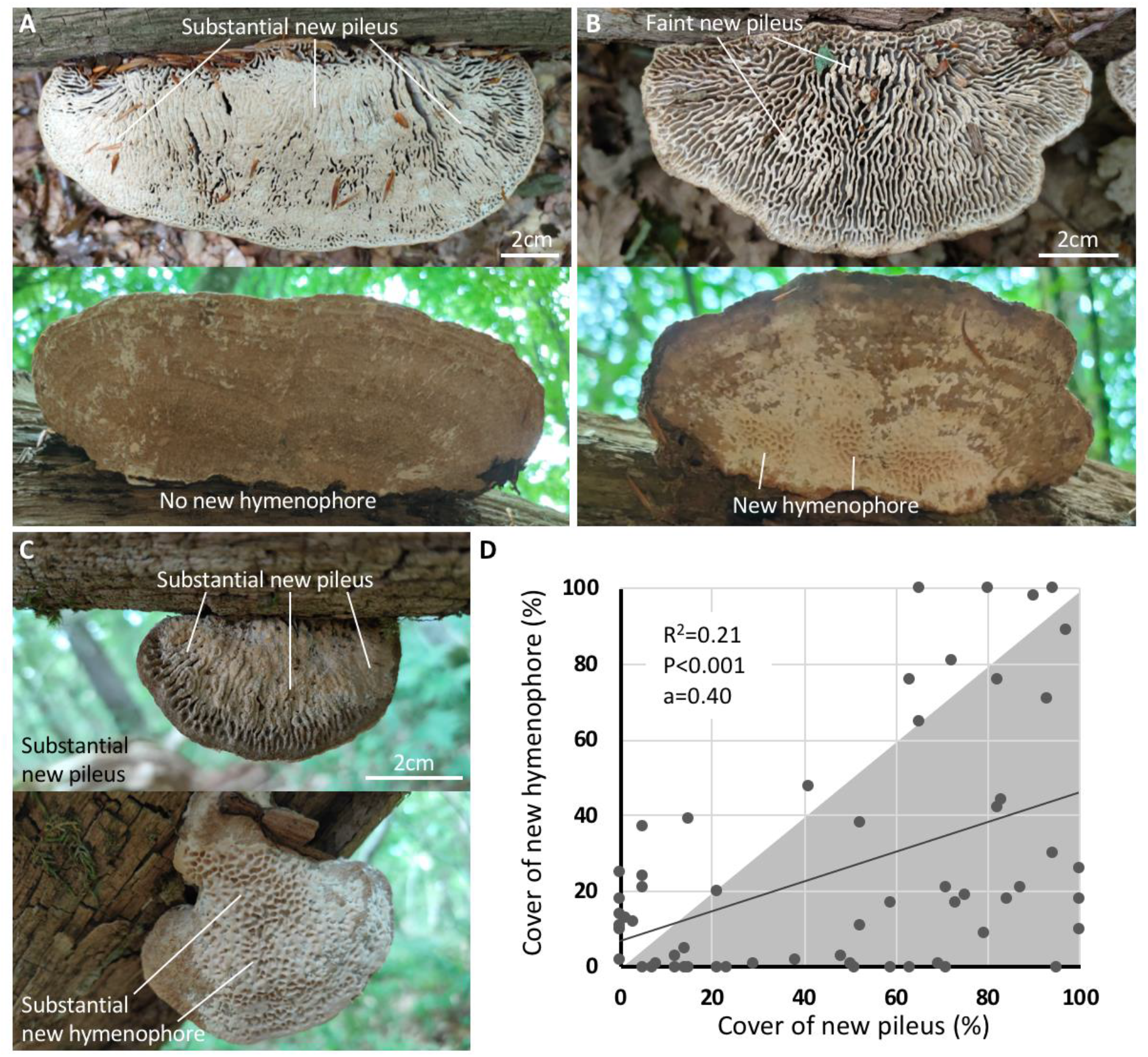
One-year changes to experimentally flipped basidiocarps. A. Example of basidiocarp that has grown a substantial new pileus, whereas there has been no growth of a new hymenophore. B. Example of basidiocarp with faint growth of a new pileus, whereas the growth of a new hymenophore is more pronounced. C. Example of basidiocarp with substantial growth of a new pileus and hymenophore. D. Pearson’s linear regression analysis (P<0.001) of experimentally flipped basidiocarps with growth response (N=59), showing that most basidiocarps had a more extensive new pileus than new hymenophore (grey triangular area).

That the new hymenophores had a poroid configuration appeared to be transitory, however. By July 2022, 25 basidiocarps exhibited a kind of ‘broken’ poroid configuration (Figure 7). This configuration approximated the one observed in the naturally flipped basidiocarps, notice for example how similar the basidiocarps are of Figures 4A and 7A. In both naturally and experimentally flipped basidiocarps, the ‘broken’ poroid configuration was confined to the vertically grown hymenophore, whereas the new hymenophore that had grown outwards was predominantly lamellate. Taken together, these observations and experiments show firstly that the new hymenophore does not appear to grow from the margin towards the base as could be expected if a marginal pore field was solely responsible for the configuration of the hymenophore. Secondly, the new hymenophore does not maintain the configuration of the old hymenophore as would be expected if the pore field patterns a fixed configuration.

**Figure 7.**
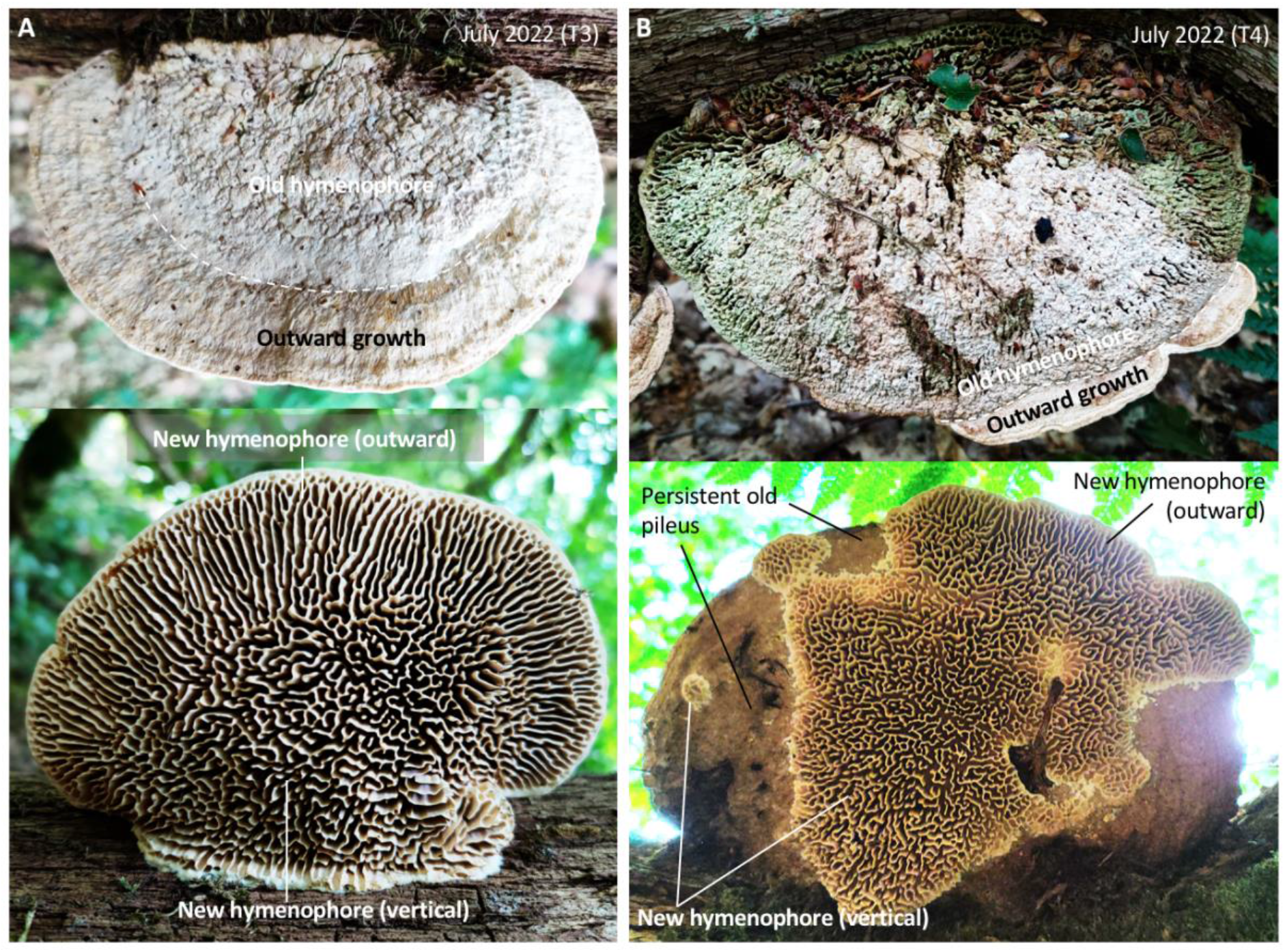
Configuration of new hymenophore two years since being experimentally flipped. A. Example of basidiocarp from Flipped trunk 3 in which the old hymenophore is entirely covered by a new pileus, the old pileus is entirely covered by a new hymenophore, and there has been outward growth. There is a broken poroid configuration to the new hymenophore that has grown vertically, whereas the outward-grown hymenophore is lamellate. B. Example of basidiocarp from Flipped trunk 4 in which the old hymenophore is not fully covered by a new pileus, the old pileus is not fully covered by a new hymenophore, and there has been outward growth. There is a broken poroid configuration to the new hymenophore that has grown vertically, whereas the outward-grown hymenophore is lamellate.

We noticed a large fraction of the studied basidiocarps were not responsive and that among the responsive basidiocarps there was much variation in the extent of new growth. A final series of experiments were made to try to reproduce a common perturbation to normal growth, namely growing around obstacles, which would give insight into whether a large fraction of unresponsive basidiocarps is to be expected and at the same time allow for the measurement of growth rates.

### Natural growth around obstacles and growth around bands

Encountering obstacles such as twigs, constitute a common perturbation to the normal growth of basidiocarps (Figure 8). Perturbations to normal growth can be revealed as indentations to the margin and altered configuration to the hymenophore (Figure 8A-A’). Greater obstacles tend to associate with greater perturbations (Figure 8B-B’). Based on the observations of 53 basidiocarps in various stages of growing around twigs of various sizes, it was deduced that the encounter of obstacles will entail growth around the obstacle rather than through it. The distance around the obstacle, or perimeter, is greater than the distance through it. Consequently, if all parts of the margin grow approximately at the same rate, the part of the margin that grows around the twig will expand around the twig and less so distally. This results in the indentation of the margin. A zone of tube apposition is formed when margins merge and this can persist over a number of growth periods. A very similar zone of apposition can be found where two basidiocarps merge (Supplementary Figure 1). If growth is sustained since the incorporation, the margin may become normal. In this scenario, all parts of the margin are growing outward at a similar rate and such growth in effect resembles fluid flow around a cylinder (e.g. Mo et al 2013).

**Figure 8.**
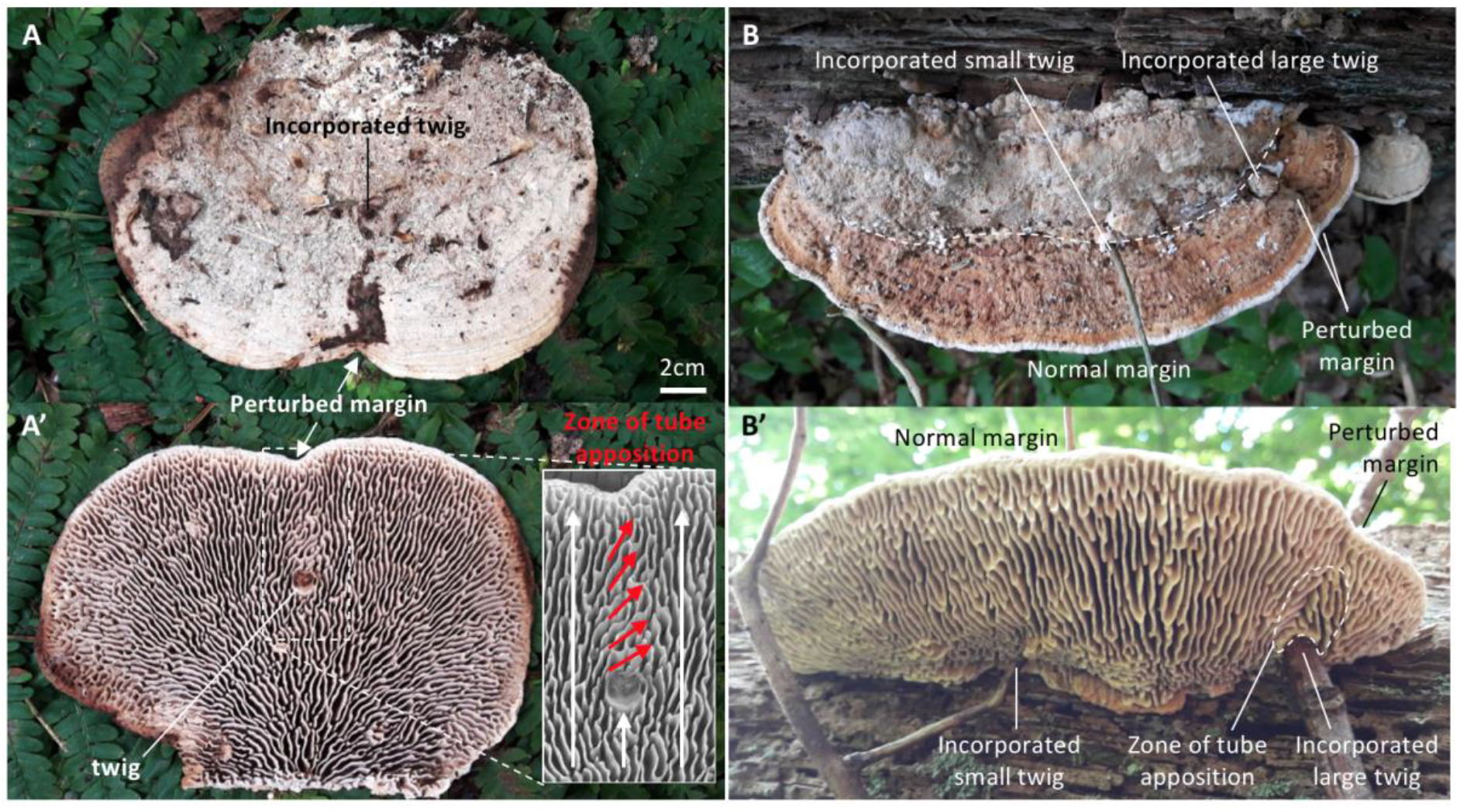
Observations on incorporation of naturally occurring obstacles. A. Basidiocarp with a single incorporated twig associating with a single indentation of the margin and a skewed orientation of the lamellate tubes between the twig and the margin. B. Basidiocarp with a small and a large twig incorporated in the same growth zone (dashed line). Only in association with the large twig is the margin (and pileus) perturbed, suggesting that a greater obstacle associates with greater perturbation.

Of the 48 experimentally banded basidiocarps, after one year 41 were still situated in the original position and had the bands attached (33 single band; 8 double band). One trunk with five banded basidiocarps had fallen over, bands had been ripped off two basidiocarps, and these amounted to the seven lost basidiocarps. Of the 41 remaining basidiocarps, 29 exhibited growth at the margin since banding (Figure 9) and 12 exhibited no growth. The growth of the 29 basidiocarps was on average 6.5 mm in the period July 2020 to July 2021 and with considerable variation between basidiocarps (range 0.3-17.7 mm, standard deviation 4.4 mm). The basidiocarp shown in Figure 9A-B grew 6.6 mm. In 19 of the 29 growing basidiocarps, the strips appeared to become partially covered by the hymenophore (Figure 9A’-B’). This shows geotrophic growth of the hymenophore and suggests that the hymenophore has a capacity to form new hymenophore at some distance from the pore field. The following year, by July 2022, four of the 29 basidiocarps had grown less than 0.5 mm and were consider to have not grown. The remaining 25 basidiocarps had on average grown 10.5 mm from July 2020 to July 2022 and with much variation between basidiocarps (range 1.0-30.2 mm, standard deviation 7.6 mm) (different growth patterns are illustrated in Figure 10). In the basidiocarp shown in Figure 9A-C, the parts that have grown since banding are similar to those of the basidiocarps that have grown around natural obstacles (Figure 8). Also, that the perturbation to growth is affected by the size of the obstacle (Figure 8B) was reproduced in the basidiocarps with bands of two different sizes (Figure 9D-D’). Taken together, this data documents that it can be expected that a large fraction of basidiocarps will not respond and that there is much variation between the basidiocarps that do respond.

**Figure 9.**
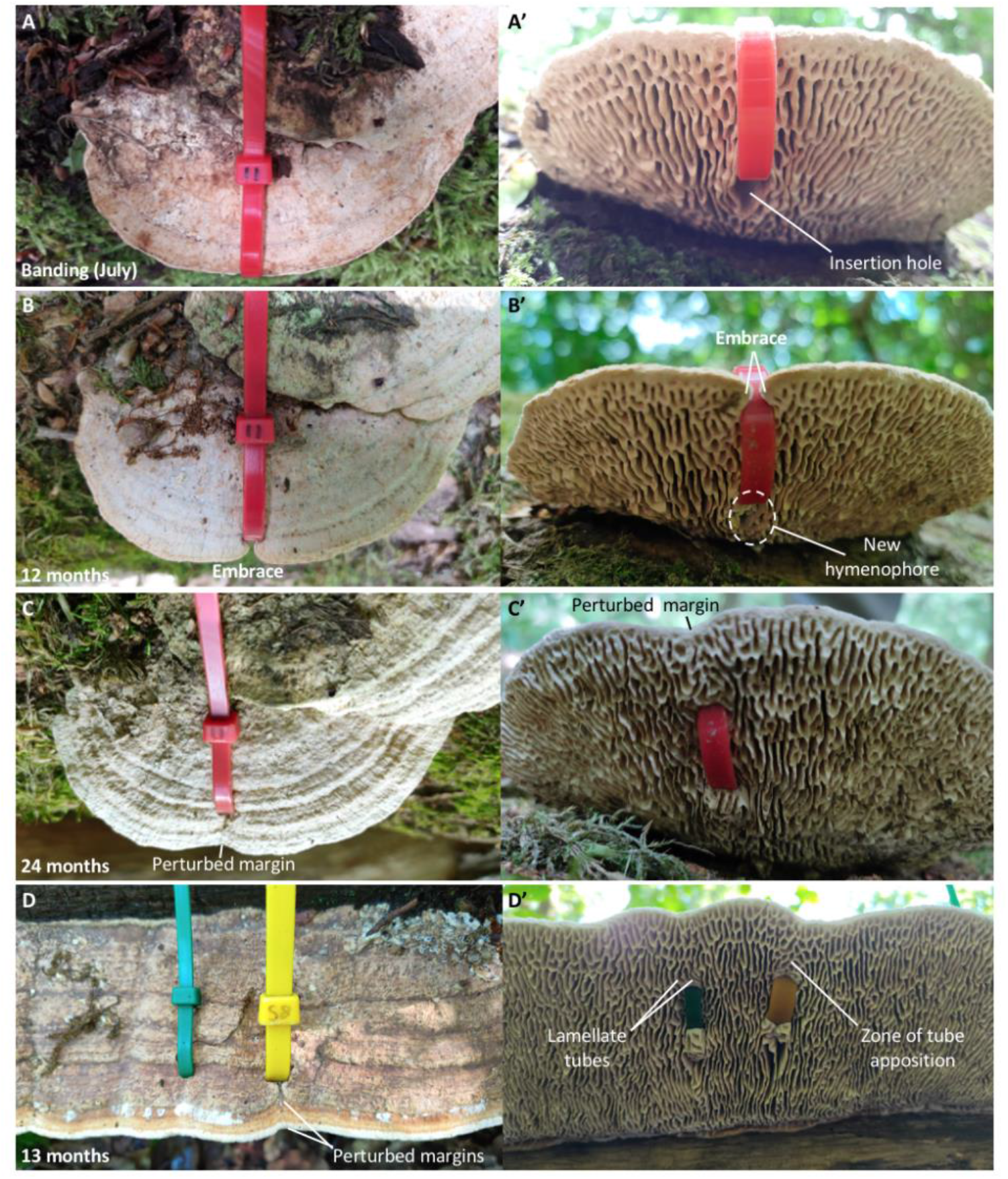
Observations on experimentally banded basidiocarps. A. Basidiocarp with attached 4.8 mm wide plastic strip (July 2020). A’. The insertion of the plastic strip resulted in a hole in the hymenophore. B-B’. One year later (July 2021), the outward growth was 6.6 mm and such that the margin now embraced the plastic strip. Notice that new hymenophore has appeared in the insertion hole, whereas the surrounding hymenophore did not change configuration. C-C’. After two years (July 2022), the outward growth was 11.6 mm but the merged margin was still perturbed. D. Basidiocarp banded with a standard-size strip (green, 4.8 mm) and a wide strip (yellow, 7.6 mm), 13 months after banding (July 2020 to August 2021). Outward growth (21.5 mm) was most perturbed distal to the wider strip. D’. Lamellate tubes were found distal to the standard-size strip, showing a poroid configuration is not the default configuration to perturbation. A zone of apposition was more evident distal to wide strip, indicating wider obstacles imposes greater perturbations.

**Figure 10.**
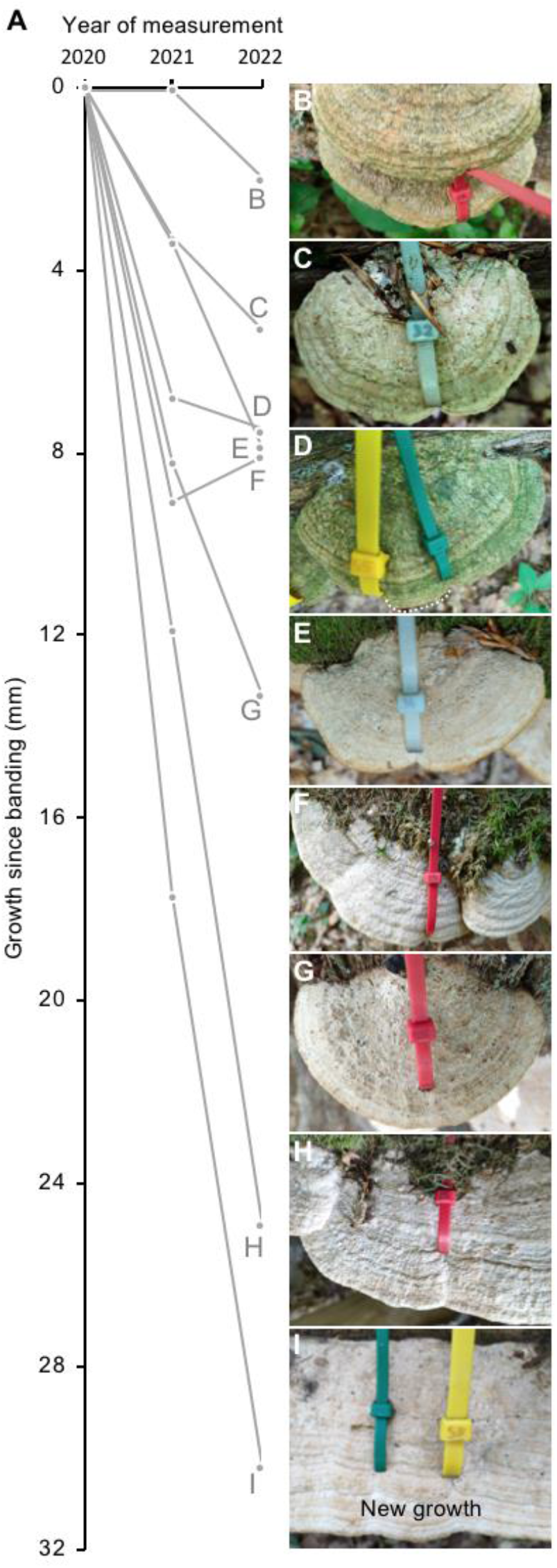
Variation in magnitude and year of growth. A. Examples of growth curves to illustrate the variation and magnitude of growth of the monitored basidiocarps. B exemplifies growth in the second year only. C and E exemplify relative slow growth, whereas D and F exemplifies little or no growth in the second year. G-H exemplify very substantial growth of basidiocarps from three different trunks. I is the basidiocarp that grew the most of all monitored basidiocarps.

## Discussion

That basidiocarps with a fully resected pore field can grow new hymenophore, and pileus, is perhaps the strongest argument against the pore field being necessary for the formation of pores. However, the regeneration of the pore field may be the first response to its resection. It is therefore interesting that basidiocarps with resected sides did not show preferential growth from the intact pore field, but rather a similar growth response across the entire cut surface, irrespective of whether the cut surface is within the hymenophore or pore field. This indicates that the capacity to configure the hymenophore resides in the basidiocarp at large. In support hereof is that the new hymenophore of flipped basidiocarps was not predominantly close to the pore field, even though the pore field was intact, but rather it appeared in spots on the pileus and not infrequently close to basidiocarp base. Also, appearance of new hymenophore was often found in the insertion hole of the banded basidiocarps, far from the largely unperturbed pore field. The data presented here, suggest the pore field does not drive hymenophore configuration. Instead, the pore field may be a manifestation of growth rather than a driver of it.

In spite of the frequent lamellate features of the oak mazegill hymenophore, the default configuration appears to be poroid since the tubes closest to the margin often take the appearance of pores (Ames 1913). Given the phylogenetic position of oak mazegill within Polyporales (Nagy et al 2016) and the presence of a mostly poroid hymenophore in other species of Daedalea (Lindner et al 2011; Han et al 2015), a poroid configuration is also the primitive condition. The data presented here on the flipped basidiocarps, show the initial appearance of geotrophically grown hymenophores is invariably poroid. Nonetheless, the poroid configuration is not the default response to perturbation since the hymenophores that grew from cut surfaces tended to be lamellate. This suggests it takes the additional factor of outward growth to form lamellate tubes. In naturally occurring and growing basidiocarps, it may be that the presence and length of lamellate tubes has a relation to growth rate and that a predominantly poroid hymenophore reflects a slowly grown basidiocarp.

The growth rate was found to be on average some 5 mm, in the period July 2020 to July 2022. This suggests that large basidiocarps from the same localities, some of which clearly exceeds a base-margin distance of 5 cm, may be more than a decade old and an age of two decades seems plausible for some basidiocarps. The growth rate varied more than an order of magnitude, between 0.5 and 15 mm per year, and if slowing-growing basidiocarps can reach large sizes, such basidiocarps can be several decades old. One caveat to the accuracy of the measured growth rates is that they are were measured on basidiocarps that had been damaged by insertion of one or two plastic strips. While it was not obvious that banded basidiocarps grew slower than the neighboring and intact basidiocarps, it was not ruled out their growth was affected by the banding.

### Conclusion

Basidiocarps of the oak mazegill do not grow fast, but they can attain a very substantial size over time. Consequently, they can reach a considerable age. Despite their growth rate, they can exhibit a pronounced response to reorientation and resection within weeks. This response includes the formation of a new hymenophore, which seems to grow from the basidiocarp at large. Therefore, it is suggested that the pore field does not drive hymenophore configuration. Perhaps the pore field is a manifestation of growth rather than a driver of it.

## Acknowledgements

Gratitude is extended to the inhabitants of Nonbo hede 6 and to Irina Sergeeva for technical support and critical reading of a draft version of the manuscript.

## Figures and figure legends

**Supplementary Figure 1.**
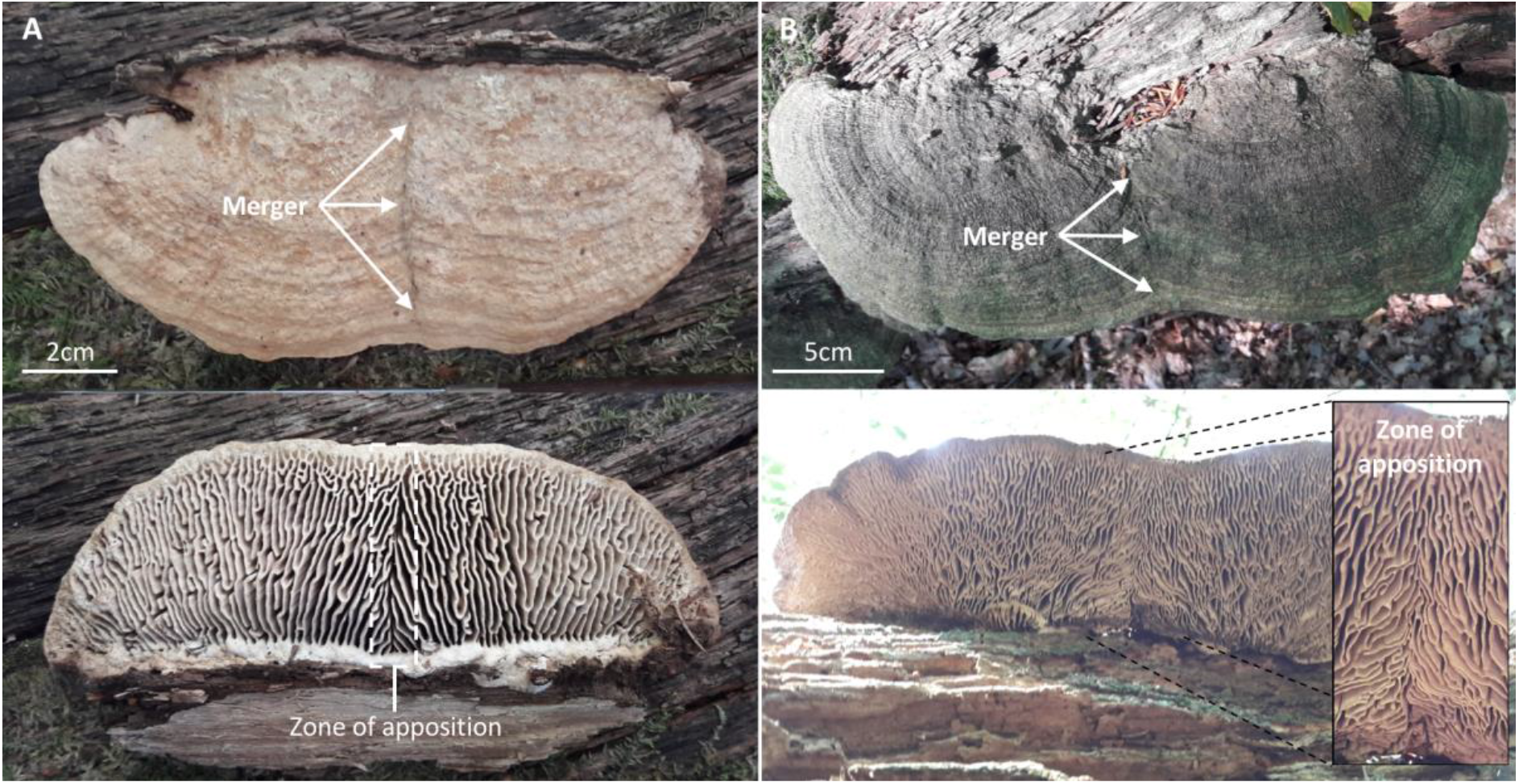
Zone of apposition of the hymenophore where 2 basidiocarps merge in a medium-sized (A) and a large (B) basidiocarp.

